# Neural signature of everyday function in older adults at-risk of cognitive impairment

**DOI:** 10.1101/2021.12.16.472826

**Authors:** Pierfilippo De Sanctis, Johanna Wagner, Sophie Molholm, John J. Foxe, Helena M. Blumen, Douwe J. Horsthuis

## Abstract

Assessment of everyday activities are central to the diagnosis of pre-dementia and dementia. Yet, little is known about the brain substrates and processes that contribute to everyday functional impairment, particularly during early stages of cognitive decline. We investigated everyday function using a complex gait task in normal older adults stratified by risk of cognitive impairment. We applied a novel EEG approach, which combines electroencephalographic with 3D-body tracking technology to measure brain-gait dynamics with millisecond precision while participants are in motion. Twenty-six participants (mean age = 74.9 years) with cognitive and everyday functional profiles within the normal range for their age and sex were ranked for risk of cognitive impairment. We used the Montreal Cognitive Assessment battery, a global index of cognition with a range from 0 to 30, to classify individuals as being at higher (22-26) and lower risk (27+). Individuals walking on a treadmill were exposed to visual perturbation designed to destabilize gait. Assuming that brain changes precede behavioral decline, we predicted that older adults increase step width to gain stability, yet the underlying neural signatures would be different for lower versus higher risk individuals. When pooling across risk groups, we found that step width increased and fronto-parietal activation shifted from transient, during swing phases, to sustained across the gait cycle during visually perturbed input. As predicted, step width increased in both groups but underlying neural signatures were different. Fronto-medial theta (3-7Hz) power of gait-related brain oscillations were increased in higher risk individuals during both perturbed and unperturbed inputs. On the other hand, left central gyri beta (13-28Hz) power was decreased in lower risk individuals, specifically during visually perturbed input. Finally, relating MoCA scores to spectral power pooled across fronto-parietal regions, we found associations between increased theta power and worse MoCA scores and between decreased beta power and better MoCA scores.Able-bodied older adults at-risk of cognitive impairment are characterized by unique neural signatures of mobility. Stronger reliance on frontomedial theta activation in at-risk individuals may reflect higher-order compensatory responses for deterioration of basic sensorimotor processes. Region and spectral-specific signatures of mobility may provide brain targets for early intervention against everyday functional decline.

## Introduction

Assessment of everyday activities are central to the diagnosis of pre-dementia and dementia^1^. Yet, little is known about the brain substrates and processes that contribute to everyday functional impairment, particularly during early stages of cognitive decline. Based on current clinical guidelines, mild problems performing instrumental daily activities such as shopping or using public transportation can occur during the stage of mild cognitive impairment (MCI) syndrome.^1,2^ It is largely unknown, whether milder problems performing everyday activities and underlying neural correlates are discernable before individuals reach the MCI stage. A better understanding of everyday function during the pre-MCI stage is important to better characterize those at risk and develop early evidence-based intervention.^3-7^

The ability to adjust gait to complexities in our environment is integral to many instrumental daily activities. Brain changes in relation to gait performance have been described using magnetic resonance imaging (MRI). Studies link gray matter volume with gait speed^8^ and variability^9^ in regions known to activate during gait such as basal ganglia, primary and supplementary motor area, and prefrontal cortex. Importantly, slower gait speed in non-demented community-dwelling older adults has been linked to poorer ability to use public transportation, do laundry, manage medication, etc.^10^ Overall, imaging findings underscore the relevance of gait-related brain changes as early signs of everyday functional limitation. An important next step is to apply light-weight portable technology to more directly investigate brain processes during actual movement in the context of everyday function.

Mobile Brain/Body Imaging (MoBI) combines electroencephalographic (EEG) with 3D body tracking technology to measure brain-gait dynamics with millisecond precision during walking. We^11-16^ and other groups^17-21^ have demonstrated feasibility of high-quality EEG recordings during walking. Several electrophysiological signatures associated with movement have been identified and two of central interest to this study are presented here. First, neuronal oscillations in the mu (8-12Hz) and beta (13-28Hz) frequency range represent activation of sensorimotor cortex.^22-28^ Mu and beta amplitude decreases are seen during preparation and execution of movement, with effector-specific (e.g., foot, finger, tongue) distributions in line with the somatotopic arrangement of pre and post central gyri. Recent MoBI studies reported phase-related mu/beta amplitude decrease coinciding with the swing phase of the gait cycle prior to foot placement.^29-31^ Interestingly, in more complex walking tasks a shift from transient to sustained mu/beta decrease across the gait cycle can be observed.^29,31,32^ Yokoyama^31^ compared walking to precision walking by asking participants to adjust each step to hit a series of visually presented ‘stepping pads’ projected onto the treadmill belt in front of them. Distance between pads varied randomly. Walking and precision walking were marked by transient mu/beta sensorimotor activation coinciding with the swing phase. When comparing both tasks directly against each other however, mu was decreased across the gait cycle indicating that precision walking led to sustained sensorimotor activation. Similar shifts were observed in tasks that required stepping in synch with an auditory pacemaker^32^ and using visual feedback to track a target speed.^29^ Overall, studies that emphasize sensory guided walking tasks show stronger activation (i.e., mu/beta suppression) of pre/post central gyri across the gait cycle. This may reflect added demands for transforming sensory input into motor output to accomplish goal-directed locomotion.^26,33-35^

The second measure of interest observed in complex mobility tasks is medial-frontal theta (3-7Hz) oscillations, which have been interpreted as a neural correlate of postural control.^11,13,36-41^ In a balance beam walking task, impending loss of balance was marked by theta increase during the transition from swing to stance phase of the gait cycle prior to stepping off the beam^19^. In our own work, we exposed participants to a large-scale visual display of dots radiating outward from a central point of extension while they walked on a treadmill.^11,13^ To destabilize individuals, dots shifted in the mediolateral direction superimposed onto the outward radiation. Compared to an unperturbed condition (i.e., static image of dots), participants walked with shorter and wider strides, which we interpreted as cautious gait. Gait changes were accompanied by power modulations in theta localized to medial-frontal area. In another study^11^, using the same stimulation but asking participants to stand with feet side-by-side or heel-to-toe, we tested neural correlates of postural control in young and cognitively normal older adults. Medial-frontal theta increased during visually perturbed input in both groups, but less so in older adults, who exhibited larger postural sway particularly during heal- to-toe standing. These and other findings^38,39^ implicate premotor theta in adjustment of posture and gait to maintain stability.

In the current study, we sought to identify neural signatures of mobility in able-bodied older adults during early stages of cognitive decline. We exposed twenty-six older adults to visual perturbations while walking. The Montreal Cognitive Assessment (MoCA) score was used to divide the sample (cut-off score ≤ 26; range 22 to 30) and as a continuous variable to rank older adults from lower to higher risk of cognitive impairment. Assuming brain changes precede decline in everyday function, we hypothesized that all participants would increase base-of-support by widening step width to gain stability during visually perturbed compared to unperturbed inputs. Yet, the underlying neural signatures would differ between individuals at lower and higher risk as measured by the MoCA. Specifically, mu/beta power over central gyri would be decreased and shifted from transient to sustained during perturbed compared to unperturbed input. This decrease and shift, however, would be less pronounce in at-risk individuals. Finally, medial-frontal theta would be increased during perturbed input – but less so in at-risk individuals.

## Materials and methods

### Participants

Twenty-six older adults (14 women, mean age = 74.9) participated in the study. Participants were cross-enrolled from the Cognitive Control of Mobility in Aging (CCMA) Study, a longitudinal cohort study probing brain predictor of mobility in aging. Details have been described previously.^42-44^ For the purpose of this study, individuals adjudicated by consensus CCMA conference as cognitively normal were asked whether they were interested to be contacted for future studies. If so, an initial phone screening was conducted to assess general health and mobility. We used the AD8, a brief phone interview, to screen (AD8 score ≤ 1) for evidence of cognitive impairment.^45,46^ Older adults were re-evaluated on site to ensure their cognitive status did not change since their last CCMA assessment. Mean time between assessments was 1.4 years. Tests included the MoCA^47^, the short form of the Geriatric Depression Scale^48^, the Timed Up and Go test^49^, the Fall Self-Efficacy Scale^50^, the Lawton Instrumental Activities of Daily Living Scale^51^, and the Trail-Making Test and Color-Word Interference Test from the Delis-Kaplan Executive Function System (D-KEFS).^52 53^ We used the MoCA to estimate risk of cognitive impairment. The MoCA is a cognitive screening test with a cut-off score of <24/30 yielding a sensitivity of 90% and specificity of 87% to differentiate mild cognitive impairment from normal^54^. In our sample we used a cutoff score of ≤ 26 to classify individuals into groups of higher (22-26) and lower risk (27-30) for cognitive impairment. To further complement phenotypic assessment of our sample, we included mobility, functional, and cognitive test scores collected as part of the CCMA study. Quantitative gait measures were obtained using a computerized walkway (180 × 35.5 × 0.25 inches) with embedded pressure sensors (GAITRite; CIR Systems, Havertown, PA)^55^. Complex everyday function were assessed using items from the disability component of the Late Life Function and Disability Instrument (LLFDI) and the Lawton Instrumental Activities of Daily Living.^51,56,57^ The Repeatable Battery for the Assessment of Neuropsychological Status (RBANS) is a standardized clinical tool measuring attention, language, visuospatial/constructional abilities, and immediate and delayed memory^58^. Studies report good discrimination for detection of mild cognitive impairment, with receiver operating characteristic analyses yielding area under the curve of 0.88 for the Total RBANS score and 0.90 for the delayed memory score. Further inclusion criteria were normal or corrected-to-normal vision, free from any neurological or psychiatric deficits or disorders likely to affect gait and able to walk on a treadmill for approximately one hour. The Institutional Review Board approved the experimental procedures which were in compliance with the Declaration of Helsinki. All participants provided written informed consent and were compensated ($15 per hour).

### Experimental Design and Procedure

The design is part of a more extended study testing effects of visual perturbation as well as performance of a secondary cognitive task on mobility.^11,13^ Here, we limit our results on the effects of visual perturbation on gait as we were specifically interested to probe neural correlates of gait adjustment (for a full description pleases see^13^). Participants walked on a treadmill (Tuff Tread® 4608PR) instructed to fixate a central fixation cross. Participants performed four visually perturbed and four unperturbed walking blocks (each about 3.5 minutes long) randomized within participants. During perturbed stimulation, a large-scale visual field of dots was projected centrally (InFocus-XS1-DLP) onto a black wall 1.5m in front of the participant. Field of view extended approximately 100° horizontally by 100° vertically. The stimulation consisted of 200 randomly placed white dots emanating outward from a central focus of expansion point. Superimposed to the outward motion was a sinusoidal perturbation in the mediolateral direction. Flow was programmed from:

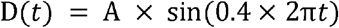

D(t) was the translation distance (meter), A was the amplitude of displacement and *t* was time (sec). Sinusoidal perturbations in the mediolateral direction were applied at amplitudes of 0.12 meter. The frequency selected (0.4 Hz) was within the range used in previous perturbation studies on gait and balance.^59,60^ Static dots placed randomly across the visual field served as control condition (i.e., no perturbation). Stimulation was programmed with Presentation (Neurobehavioral Systems, Berkeley, CA). Participants familiarized themselves with the walking task before undertaking the main experiment. To minimize fatigue participants took a short break after each block. A self-selected walking speed was chosen at the beginning of the experiment and maintained throughout. All participants walked while wearing a safety harness.

### Kinematics recordings

Optitrack infrared motion capture system with 9-cameras was used to collect kinematic data in the X/Y/Z direction at a sample frequency of 100Hz (Arena v.1.5 acquisition software, Natural Point). We placed three markers on each foot over the participants’ shoes: on the calcanei, the second and the fifth distal metatarsals.

### Electrophysiological recordings

Continuous EEG was recorded with a 160-channel BioSemi ActiveTwo system (digitized at 512Hz; 0.05 to 100 Hz pass-band, 24 dB/octave). Time-synchronized acquisition of EEG and rigid body motion tracking was conducted with Lab Streaming Layer software (Swartz Center for Computational Neuroscience, UCSD, available at: https://github.com/sccn/labstreaminglayer).

## Data Analysis

All EEG and kinematic data analyses were performed using custom MATLAB scripts (MathWorks, Natick, MA) and EEGLAB (Version 2019.1).^61^

### Gait Kinematics

Spatiotemporal features of gait kinematics were determined in the following manner: We equated kinematic and electrophysiological time series by up-sampling kinematic data from 100 to 512Hz. Kinematic data were then lowpass filtered using an IIR filter (Butterworth order two) to remove high-frequency artifacts and highpass filtered at 0.1Hz (Butterworth Filter order 4) to remove slow drifts. The filtered position time-courses were then averaged across reflective markers at each foot for the horizontal (anterior-posterior) direction and for the vertical direction (up-down). For these four averaged position time-courses anterior-posterior and up-down for left and right foot, we then computed the velocity profile. We then estimated the toe-off from the velocity profile of the up-down position and the heel-strike from the velocity profile of the anterior-posterior position.

### EEG Preprocessing

EEG preprocessing was followed by an Independent Component Analysis (ICA)^62^ and dipole-fitting approach^63,64^, similar to methods reported in previous MoBI studies.^17,20,32,36,65,66^ Integrated analysis of gait kinematic and electrophysiological data was performed using custom MATLAB scripts (MathWorks R2018b, Natick, MA) and EEGLAB. Continuous raw data were re-referenced to CPz, high pass filtered at 2 Hz (zerophase FIR filter order 3380) and concatenated across blocks. An automatic channel rejection procedure^67^ was applied to exclude channels with flat lines, correlation between neighboring channels < 0.6, and values of line noise exceeding signal by 8 standard deviations. Data were visually inspected and additional channels were excluded if artefacts were present over ∼50 sec. On average, 3.9 EEG channels were removed for further analysis (range: 0-17). Data were re-referenced to average reference. Next, an adaptive independent component analysis mixture model algorithm (RUNICA) was used to decompose EEG signals into independent components (ICs). Resulting ICs were co-registered with a standard Montreal Neurological Institute boundary element head model and fit with single equivalent current dipole models using the DIPFIT toolbox in EEGLAB.^63,64^ Only dipoles located within the brain, with a fit accounting for at least 80% of the variance for a given IC scalp projection, were retained.^68^ In addition, ICs originating from eye blinks, bad electrodes, cable sway and muscle activity noise were rejected.^69^ The average number of brain ICs included for further analysis was 6.4 (ranging from 1 to 11).

### Cortical IC Clustering

Brain ICs were clustered across subjects using feature vectors coding for IC difference in power spectral density, dipole location, and scalp projection.^18,20^ Feature vectors were reduced to 10 principal components and clustered across subjects using k-means. ICs greater than three standard deviations from a cluster centroid were classified as outliers. Cluster centroids localized to occipital, bilateral sensorimotor, left posterior parietal, and medial-frontal premotor areas. Regions of interest detailed in our predictions included sensorimotor and frontal regions. Therefore, posterior parietal and occipital clusters were excluded from further analysis. Table 1 lists the numbers of ICs within each cluster and the cluster centroid locations.

**Table 1.**
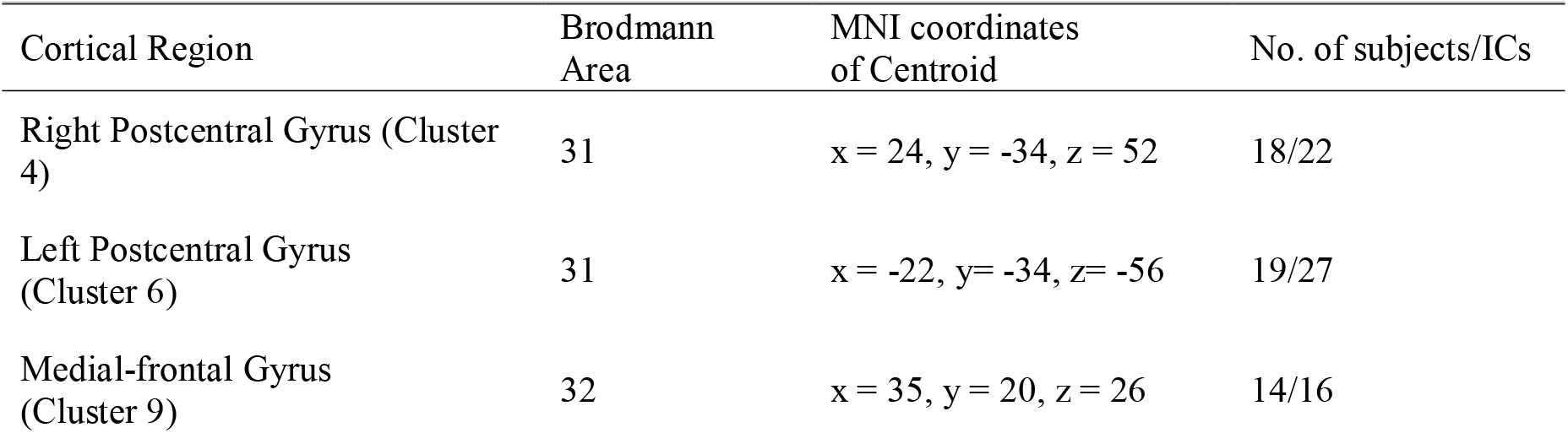
Source localized cluster. Approximate location (Brodmann area and Talairach coordinates) and descriptive information (number of subjects and independent components (ICs) included from young and old groups) for each cluster of cortical sources.

| Cortical Region | Brodmann Area | MNI coordinates of Centroid | No. of subjects/ICs |
| --- | --- | --- | --- |
| Right Postcentral Gyrus (Cluster 4) | 31 | x = 24, y = -34, z = 52 | 18/22 |
| Left Postcentral Gyrus (Cluster 6) | 31 | x = -22, y = -34, z = -56 | 19/27 |
| Medial-frontal Gyrus (Cluster 9) | 32 | x = 35, y = 20, z = 26 | 14/16 |

### Event-Related Spectral Perturbations (ERSP)

ERSP were computed from single trial spectrograms for each IC, time-locked to the right heel-strike.^*70*^ IC activity was epoched 1 second before and 3 seconds after right heel-strike. ERSPs were computed by determining the power spectra over a sliding latency window and normalizing the spectrogram by their respective mean spectra (averaged across latency window).^70^ To compute gait cycle ERSPs, we time-warped single trial spectrograms applying a linear interpolation function to align left toe off, left heel strike, right toe off, and right heel strike across epochs following the methods introduced by Gwin and colleagues^20^. Relative change in spectral power was obtained by subtracting the average log spectrogram across gait cycles from each single-trial log spectrogram. We computed grand average ERSPs for each condition and cluster.

## Statistical Analysis

### Gait

Repeated measures analysis of variance (ANOVA) with factors Perturbation (perturbed versus unperturbed) and Group (higher versus lower risk) was performed to determine differences in step width and step width variability.

### Grand Mean Event-Related Spectral Perturbations 3-40Hz

To test group-level activation of cortical regions, we averaged across single-subject spectrograms of ICs with cluster centroids in left/right sensorimotor and medial-frontal regions separate for perturbed and unperturbed inputs. Significance of power deviations from baseline, the mean log spectral power average across the gait cycle, were computed using bootstrapping method.^61^ ERSPs were masked for significant differences (p < 0.05) and false discovery rate (FDR) was applied to correct for multiple comparisons.^71^

### Relationship between MoCA and theta (3-7Hz), mu (8-12Hz), and beta (13-28Hz) power

Associations between MoCA and spectral EEG power were analyzed in two ways. First, we used MoCA as a continuous variable. Associations between MoCA and spectral power pooled across brain regions separate for walking with and without perturbations were tested with partial correlations adjusted for age. We also using a MoCA cut-off score of ≤ 26 to classify participants into groups of lower and higher risk for cognitive impairment and performed a 2 × 2 × 7 analysis of variance with factor Group, Perturbation, and Gait Cycle Phase. The gait cycle was divided into stance phase (0–60%) and swing phase (60–100%) according to the definition by Perry.^72^ The stance phase is further divided into: Loading response (0–10%), mid stance (10–30%), terminal stance (30– 50%) and pre-swing (50–60%). For the swing phase, there are initial swing (60–73%), mid-swing (73–87%) and terminal swing (87–100%). Separate ANOVAs for brain regions (left/right sensorimotor and medial-frontal) and spectral power band (theta, mu, and beta) were performed. Greenhouse–Geisser corrections where applied in case assumption of sphericity was violated. Post-hoc tests were computed using simple paired t-tests, controlling for false discovery rate^71^ with a significance level at 0.05.

## Data availability

The data of this study are available from the corresponding author upon reasonable request. The datasets are not publicly available as they contain information that could breach privacy of research participants.

## Results

### Demographics, functional and cognitive assessment

Table 2 lists assessment scores collected as part of the parent study and electrophysiological EEG investigation. While all scores fell within the normal range for age and sex, there were differences in some assessments between groups. With regard to measures collected as part of the EEG study, individuals at higher risk reported more concerns about falling and had lower scores on the Trail Making Test Part B. For tests obtained as part on the parent study lower scores in delayed memory are reported for individuals at higher risk. Participants did not differ significantly in terms of age, education, gender, BMI and depression scores.

**Table 2.** Participant characteristics, Means (SDs)

| | | Lower-risk CI:<br>MoCA ≥ 27 | Higher-risk CI:<br>MoCA ≤ 26 | Stats |
| --- | --- | --- | --- | --- |
| Characteristics: mean (SD) | N = 26 | n = 16 | n = 10 |  |
| MoCA | 27.2 (1.9) | 28.38(1.1) | 25.3 (1.3) | >0.001 |
| Age, y | 74.8 (4.54) | 74.9 (4.6) | 74.7 (5.3) | 0.39 |
| Gender (f/m) | 14/12 | 11/5 | 3/7 | 0.054 |
| Ethnicity: non-Hispanic Black, % | 6 | 4 | 2 |  |
| Ethnicity: Non-Hispanic White, % | 17 | 10 | 7 |  |
| Ethnicity: Hispanic,% | 3 | 2 | 1 |  |
| BMI | 26.98 (4.6) | 26.96 (4.9) | 27.35 (4.5) | 0.9 |
| Geriatric Depression Scale | 0.5 (1.02) | 0.81 (1.2) | 0.2 (0.4) | 0.14 |
| Education, y | 16.6 (2.3) | 17.1 (2.1) | 15.6 (2.7) | 0.13 |
| Treadmill Speed | 1.45 (0.25) | 1.53 (0.27) | 1.37 (0.18) | 0.11 |
| Assessments as part of MoBI data collection: May 2018 – July 2019 |  |  |  |  |
| Function |  |  |  |  |
| AD8, ranging from 0 to 8 | 0.12 (0.3) | 0.13 (0.34) | 0.1 (0.31) | 0.8 |
| Time Unipedal Stance, eyes open, sec. | 20.47 (12.02) | 23.8 (10.4) | 15.1 (13.01) | 0.07 |
| Get Up & Go, sec. | 9.3 (1.94) | 9.3 (2.2) | 9.3 (1.5) | 0.9 |
| Falls Self-Efficacy Scale | 19.2 (2.8) | 20.01 (3.09) | 17.9 (1.8) | 0.04 |
| IADL | 7.96 (0.2) | 8 (0.0) | 7.9 (0.3) | 0.24 |
| Cognition (D-KEFS) |  |  |  |  |
| Trail Making Test Number-Letter Switching | 10.8 (3.2) | 11.9 (1.) | 9.1 (4.3) | 0.02 |
| Color Word Interference Test | 10.2 (2.4) | 10.2 (2.6) | 10.2 (2.4) | 0.18 |
| Digit-Symbol Substitution Test | 10.4 (1.6) | 10.7 (1.64) | 9.9 (1.5) | 0.22 |
| Word Reading % | 87.1 (13.6) | 90.4 (11.7) | 82.02 (15.5) | 0.36 |
| CCMA Assessments: January 2017- September 2018 |  |  |  |  |
| Function and gait performance |  |  |  |  |
| LLFDI total | 69.3 (4.1) | 69.1 (3.9) | 69.6 (4.3) | 0.8 |
| Velocity | 105.7 (20.2) | 104.6 (21.9) | 106.9 (18.6) | 0.78 |
| Cadence | 102.9 (11.8) | 101.7 (13.5) | 104.1 (10.1) | 0.65 |
| Swing time | 0.42 (0.03) | 0.42 (0.06) | 0.42 (0.01) | 0.9 |
| Stance time | 0.78 (0.09) | 0.79 (0.13) | 0.78 (0.06) | 0.88 |
| Stride length | 126.3 (13.4) | 123.1 (17.5) | 129.6 (9.3) | 0.44 |
| Support base | 11.01 (3.1) | 10.1 (4.01) | 11.91 (2.3) | 0.36 |
| Cognition |  |  |  |  |
| RBANS: Immediate Memory | 100.7 (9) | 102.4 (10.1) | 99.01 (7.9) | 0.4 |
| RBANS: Delayed Memory | 96.8 (8.9) | 102.7 (11.3) | 91.01 (6.5) | 0.013 |
| RBANS: Visual Spatial | 95.9 (9.8) | 98.5 (10.2) | 93.4 (9.5) | 0.25 |
| RBANS: Language | 93.6 (7.8) | 95.5 (9.8) | 91.7 (5.9) | 0.33 |
| RBANS: Attention | 106.4 (10.3) | 108.4 (8.5) | 104.4 (12.1) | 0.3 |
| RBANS: total | 97.3 (11.9) | 102.9 (11.9) | 91.8 (11.9) | 0.02 |

#### Gait

Step width (SW) and step width variability (SWV) are presented in Figure 2. Analysis of variance revealed a main effect of Perturbation (F_1,24_ = 5.37, p =.029, □_p_^2^ = .17). No differences for step width variability between Group and Perturbation were noted.

**Figure 1.**
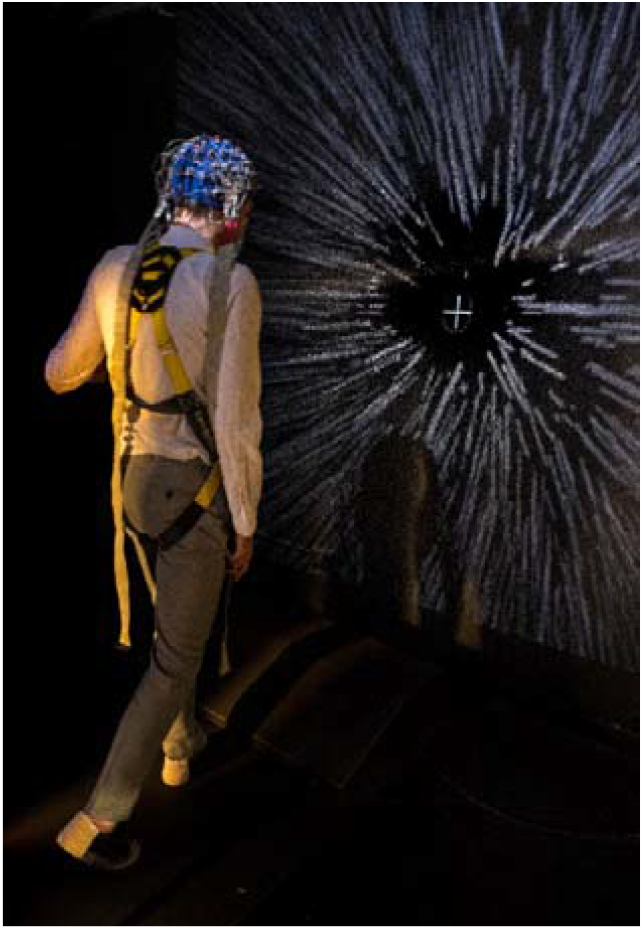
Mobile Brain\Body Imaging setup. A participant is shown wearing a 160-channel EEG cap and a safety harness while walking exposed to large-field display of dots radiation outward from the center of extension.

**Figure 2.**
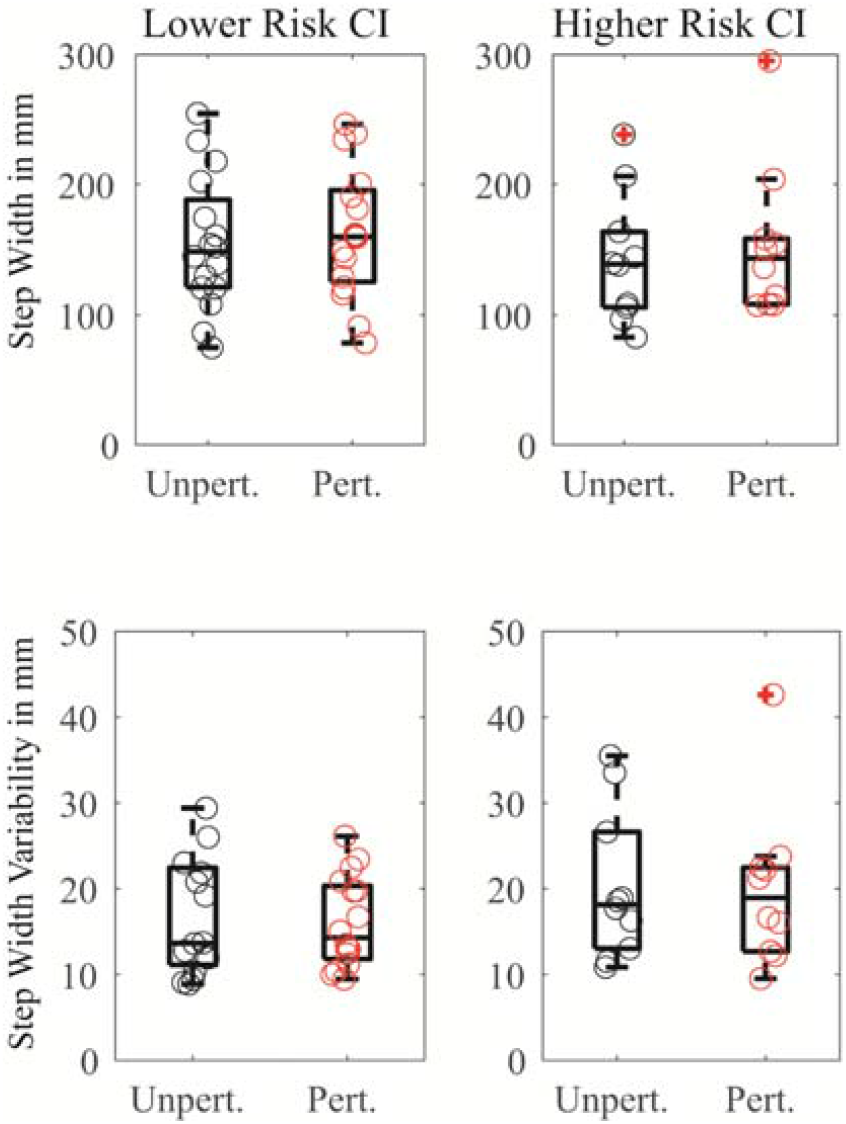
Quantitative gait markers. Step width and step width variability in participants at lower and higher risk of cognitive impairment during visually perturbed and unperturbed stimulation. Step width increased during perturbed stimulation in both groups. Crossed mark data points as outliers.

### Grand Mean Event-Related Spectral Perturbations 3-40Hz

Figure 3 shows clusters of ICs pooled across groups with centroids in or the vicinity of left sensorimotor, right sensorimotor and medial premotor regions. Group-level activation time-locked to the right heel-strike during walking with and without visual perturbation as well as differential activation were observed. Significant spectral power fluctuations were assessed relative to a common baseline: mean spectral power across the gait cycle during visually unperturbed input. Plots are FDR corrected for multiple comparisons. The main finding illustrated in Figure 3 is a shift from transient fluctuation to sustained left sensorimotor mu and right sensorimotor beta suppression across the gait cycle.

**Figure 3.**
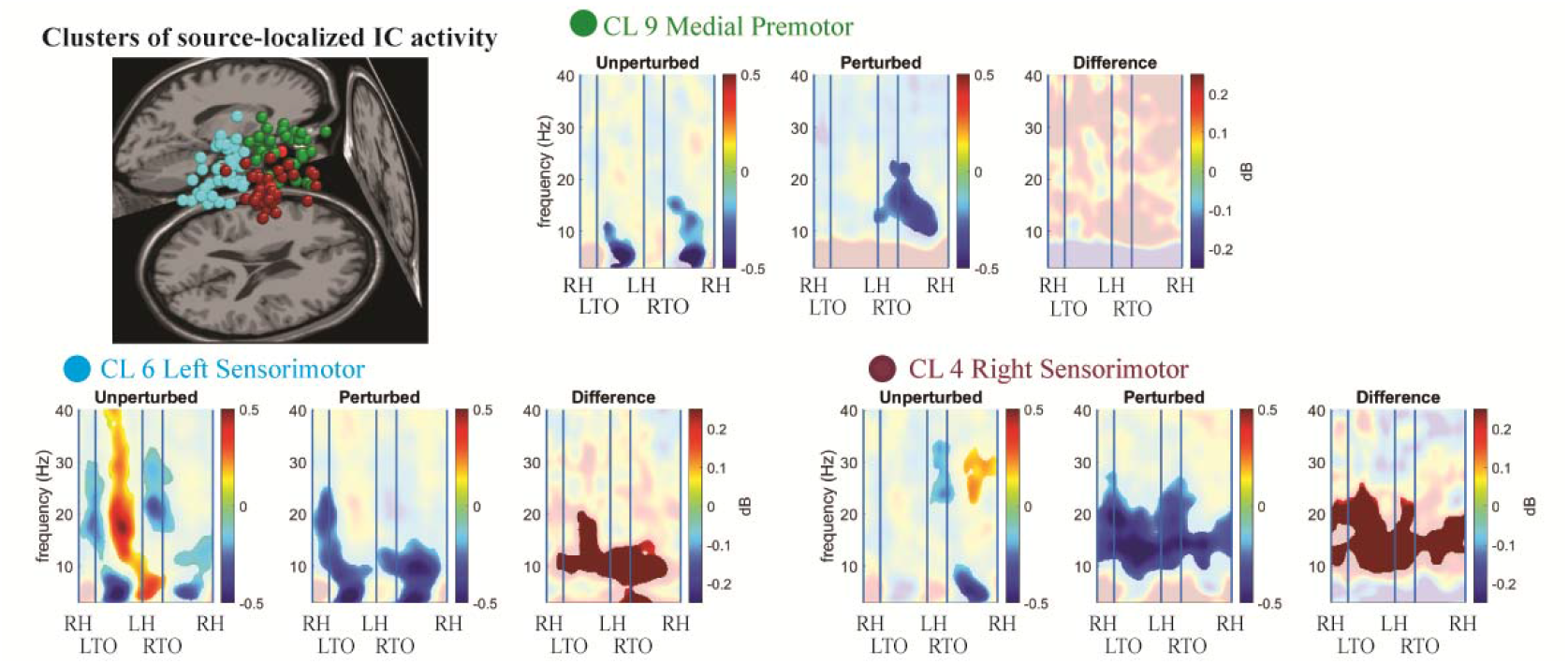
Heel-strike related cortical activation localized to frontomedial and bilateral central gyri. Group-level (across risk of CI) spectral power ratios (in dB) relative to the mean power spectrum across the gait cycle during visually perturbed and unperturbed stimulation. Transient suppression of mu and beta power during swing phases is shifted to sustained suppression across the gait cycle during exposure to perturbed stimulation. Opaque regions mask non-significant differences. Plots are corrected for multiple comparisons using FDR.

### Relationship between Event-Related Spectral Power and MoCA

ANOVA results with individuals divided into groups of lower and higher risk for cognitive impairment using a MoCA cutoff of ≤ 26 are listed in Table 3. Figure 4 illustrates mean power fluctuation and standard error of theta, mu and beta oscillations separate for the lower and higher risk group. Correlation between spectral power in theta, mu, and beta bands pooled across brain regions and MoCA score are listed in Table 2b and illustrated in Figure 5.

**Table 3.** Two by two by seven ANOVA – significant main and interaction effects in bold.

|  |  |  |  |
| --- | --- | --- | --- |
| **Brain Region** | **Frequency** | **Main and Interaction effects** | **P-value, effect size** |
| **Premotor (CL9)** | Theta | Gait Phase | $p < .001, \eta_p^2 = .33$ |
| | | Group | $p = .01, \eta_p^2 = .45$ |
| | | Perturbation | $p = .12$ |
| | | Group by Perturbation | $p = .1, \eta_p^2 = .23$ |
| | Mu | Gait Phase | $p = .028, \eta_p^2 = .18$ |
| | | Group | $p = .038, \eta_p^2 = .33$ |
| | | Perturbation | $p = .53$ |
| | | Interactions | All $p$'s $> .5$ |
| | Beta | Gait Phase | $p = .06, \eta_p^2 = .16$ |
| | | Group | $p = .54$ |
| | | Perturbation | $p = .38$ |
| | | Interaction | All $p$'s $> .4$ |
| **Left Sensorimotor (CL6)** | Theta | Gait Phase | $p = .002, \eta_p^2 = .17$ |
| | | Group | $p = .71$ |
| | | Perturbation | $p = .07, \eta_p^2 = .17$ |
| | | Group by Perturbation | $p = .06, \eta_p^2 = .18$ |
| | Mu | Gait Phase | $p = .001, \eta_p^2 = .19$ |
| | | Group | $p = .41$ |
| | | Perturbation | $p = .02, \eta_p^2 = .25$ |
| | | Interaction | All $p$'s $> .29$ |
| | Beta | Gait Phase | $p < .001, \eta_p^2 = .21$ |
| | | Group | $p = .08, \eta_p^2 = .16$ |
| | | Perturbation | $p = .33$ |
| | | Group by Perturbation | $p = .04, \eta_p^2 = .16$ |
| **Right Sensorimotor (CL4)** | Theta | Gait Phase | $p < .001, \eta_p^2 = .29$ |
| | | Group | $p = .62$ |
| | | Perturbation | $p = .58$ |
| | | Group by Gait Phase | $p = .004, \eta_p^2 = .18$ |
| | Mu | Gait Phase | $p = .04, \eta_p^2 = .12$ |
| | | Group | $p = .53$ |
| | | Perturbation | $p = .25$ |
| | | Interactions | All $p$'s $> .4$ |
| | Beta | Gait Phase | $p = .09, \eta_p^2 = .1$ |
| | | Group | $p = .22$ |
| | | Perturbation | $p = .004, \eta_p^2 = .41$ |
| | | Interactions | All $p$'s $> .3$ |

**Figure 4.**
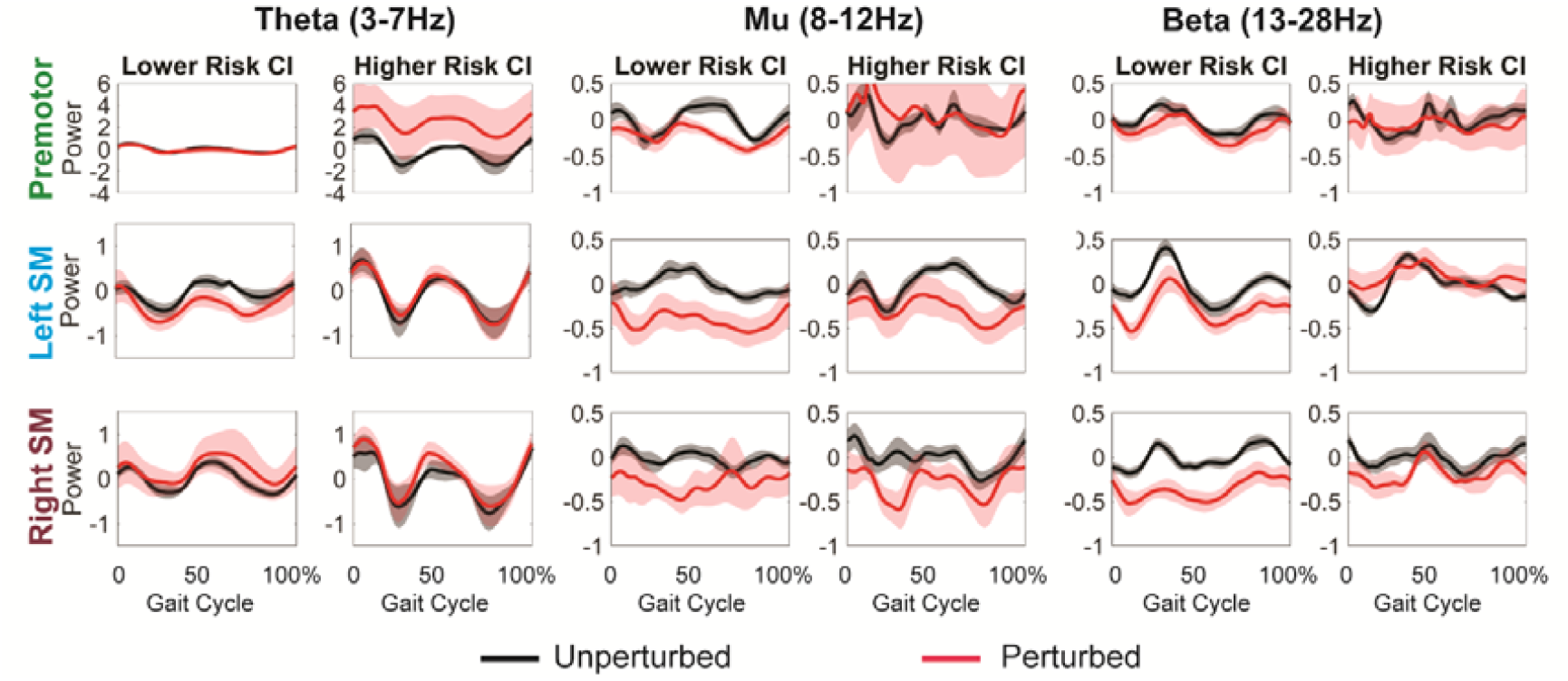
Gait-related neural oscillations in theta, mu, and beta band. Frequency-of-interest analysis with participants classified as scoring high and low (cutoff MoCA ≤ 26) within the normal range of global cognitive function. Theta increase is driven by the MoCA low-scoring group with differences between walking with and without visual perturbation seen specifically for the premotor region. Relative power decrease in mu/beta during exposure to visually perturbed input over left and right sensorimotor regions is driven mostly by the MoCA high-scoring group

**Figure 5.**
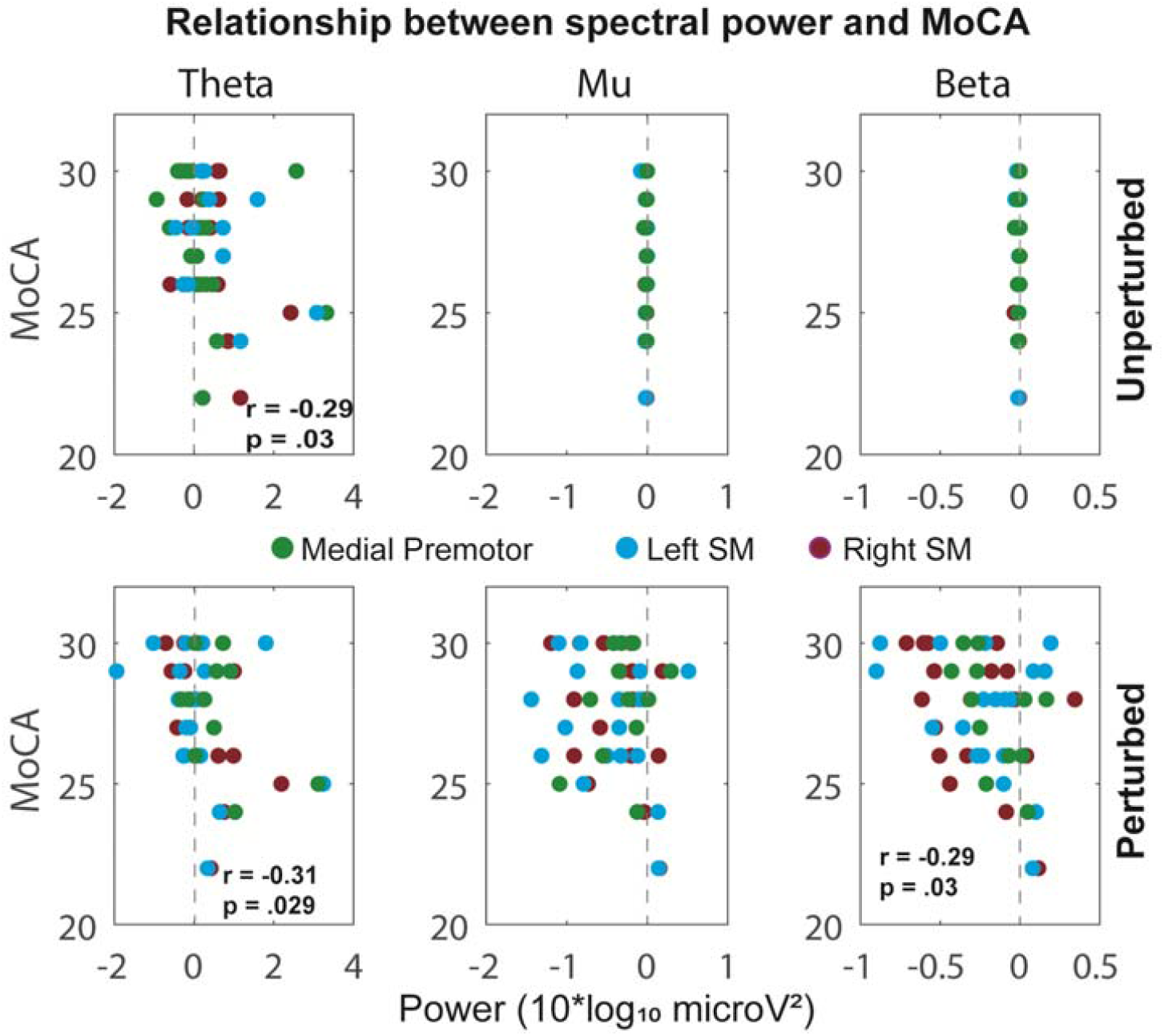
Relationships between spectral power and MoCA. Theta (3-7Hz), mu (8-12Hz), and beta (13-28Hz) power during walking with and without visually perturbed input and MoCA score. Data are pooled across brain regions. Increase in theta power during visually perturbed and unperturbed input is associated with lower scores on MoCA. Furthermore, decrease in right sensorimotor beta power during visually perturbed input is associated with higher scores on MoCA.

### Theta/dichotomized MoCA

Across brain regions, main effects for Gait Phase were observed (Right SM: F_6,96_=6.4, p <.001, □_p_^2^=.29; Left SM: F_6,108_=4.14, p=.002, □_p_^2^=.17; Medial Premotor: F_6,66_=5.39, p<.001, □_p_^2^=.33). Post-hoc tests indicate significant fluctuations coinciding approximately with double support phases of the gait cycle (Right SM: p=.05 at 0-10% & p=.04 at 50-60% of the gait cycle; Left SM: p=.015 at 0-10% & p=.08 at 50-60% of the gait cycle). Effects did not survive after Bonferroni correction. A Group effect (F_1,11_=9.27, p=.01, □_p_^2^=.45) for the medial-frontal region indicated increased theta power in the higher risk group. Furthermore, a Group by Gait Phase interaction (F_6,96_=3.4, p=.004, □_p_^2^=.18) for the right sensorimotor region indicated increased phased-related theta power in the higher risk group. Post-hoc tests did not reach level of significance during any gait phase (p’s>.13).

### Theta/continuous MoCA

Age-corrected partial correlations indicated higher spectral theta power pooled across brain regions being associated with lower MoCA score during walking with (r_partial_=-0.31, p=.029) and without (r_partial_=-0.29, p=.03) exposure to visually perturbed input.

### Mu/dichotomized MoCA

Across brain regions, main effects for Gait Phase were observed (Right SM: F_6,96_=2.2, p<.04, □_p_^2^=.12; Left SM: F_6,108_=4.2, p=.001, □_p_^2^=.19; Medial Premotor: F_6,66_=2.5, p=.028, □_p_^2^=.19). Post-hoc comparisons for the Left SM mostly survived correction and indicated stronger mu power suppression coinciding with stance and swing phases of the gait cycle (Left SM: p=.005 at 10-30%, p=.01 at 60-67%, p<.001 at 73-100%). For the medial premotor region, a main effect for Group indicated stronger mu suppression in lower risk individuals (F_1,11_=5.5, p=.038, □_p_^2^=.33). For the left sensorimotor region, a main effect for Perturbation (F_1,18_=6.1, p=.024, □_p_^2^=.25) was observed, indicating stronger mu power suppression during visually perturbed input.

### Mu/continuous MoCA

There were no significant associations between spectral mu power and MoCA score.

### Beta/dichotomized MoCA

For the right sensorimotor region, a main effect for Perturbation was observed (F_1,16_=11.01, p=.004, □_p_^2^=.41), indicating that exposure to perturbed input lead to stronger beta power suppression. For the left sensorimotor region, a main effect for Gait Phase was observed (F_1,108_=4.8, p<.001, □_p_^2^ = .21). Post-hoc comparisons suggested stronger beta power suppression during loading response (p=.002 at 0-10%) and swing phase (p=.01 at 60-73% & p=.04 at 87-100%). Comparisons did not survive correction, except for the difference noted during the loading response. Finally, stronger beta suppression in lower risk individuals during exposure to perturbed input was observed (F_1,108_=3.56, p=.04, □_p_^2^=.165). No significant beta fluctuation for the medial premotor region was observed.

### Beta/continuous MoCA

There was a significant correlation between spectral beta power and MoCA score during exposure to perturbed visual input (r_partial_=-0.29, p=.03), indicating that greater suppression of beta activity during exposure is associated with higher scores on the MoCA.

## Discussion

We set out to determine whether neural signatures of mobility among cognitively-healthy older adults are different depending on their risk of cognitive impairment. Participants in the lower and higher risk group performing a destabilizing walking task both increased step width, however the underlying neural signatures as predicted were different. Individuals at higher risk for cognitive impairment amplified fronto-medial and right central theta. In contrast, brain responses in lower risk individuals were specific to visually perturbed input and characterized by premotor mu and left central beta suppression. Furthermore, higher theta power was related to lower scores on the MoCA during both perturbed and unperturbed input. And stronger beta power suppression was related to higher scores on the MoCA. Our findings point to region- and spectral-specific differences in fronto-parietal activation underlying gait adjustment between individuals at lower and higher risk of cognitive impairment.

Substantial work established poor gait performance as an early feature of dementia.^73-85^ Our study is among the first to provide precise tracking of gait-brain dynamics in older adults^14,86,87^ to quantity moment-by-moment changes in distributed network activation involved in joint sensory, cognitive and motor processes required to orchestrate complex behaviors such as gait adjustments. Mu and beta reliance in lower risk individuals specifically during visual perturbation is in contrast to theta reliance in higher risk individuals during walking with and without exposure to visual perturbation. Our findings align well with the frequently reported posterior-to-anterior shift of brain function in aging and theories explaining such shifts towards frontal brain regions as compensatory.^88-91^ Central gyri mu/beta power suppression is a well-established neurophysiological marker of sensorimotor activation^24^ and failure to activate during visual perturbation in at-risk individuals may suggest deterioration of basic sensorimotor processes such as prioritizing inputs from visual, somatosensory, and vestibular systems based on reliability to build accurate proprioception sub-serving motor output. On the other hand, reliance on theta over the mid-frontal cortex, a region implicated in cognitive control^92,93^, may reflect higher-order compensatory responses for deterioration of basic sensorimotor processes. We acknowledge that further tests are required to firm up our interpretation in terms of compensation. For example, showing that central gyri mu/beta power suppression between standing and walking is similar and only after motor demands are further increased (i.e., exposure to visual perturbations) do individuals at higher risk manifest frontomedial theta reliance would lend stronger support to an interpretation in terms of compensation.

The strength of our study include the ecological validity of MoBI as an experimental approach to observe changes in gait and mobility-related everyday activities. We believe that imaging of gait-brain dynamics with millisecond precision in combination with the high ecological validity of the design may afford the sensitivity needed to differentiate normal from disease-related change in mobility-related everyday function during early stages of the dementia continuum.

There are several limitations to our study. Participants MoCA scores ranged from 22 to 30. Optimal cut-offs to most accurately detect MCI are debatable and recent large-scale studies with ethnically diverse populations point to different optimal cut-offs for different ethnicities.^81^ Therefore, one may question if our choice, a MoCA of 22+, is effective to screening out individuals with MCI. 25 of our 26 participants scored ≥ 24. To increase the confidence in our results, we therefore repeated our analyses excluding the person who scored 22, confirming the same pattern of results. Furthermore, interpretation of differences between groups in gait and EEG activities may be undermined by unequal numbers of participants in the lower (n=16) and higher risk (n=10) group. However, in support of our interpretation are results using the MoCA as a continuous variable and showing relationships between beta power in lower risk and theta power in higher risk individuals with MoCA scores. Also, the at-risk group showed a larger theta signal, suggesting that differences in statistical power due to difference in sample sizes are in this case not at issue.

Our findings reveal a unique neural signature of mobility-related everyday function in normal individuals at-risk for cognitive impairment. Establishing such signatures will help determine clinically relevant biomarkers in the earliest stages of cognitive decline,^94^ provide targets for non-invasive brain stimulation intervention, and further refine current criteria for prodromal or preclinical stages of dementia.

## Abbreviations

MoCA: Montreal Cognitive Assessment
MCI: Mild Cognitive Impairment
MoBI: Mobile Brain/Body Imaging
3D: Three Dimensional
Hz: Hertz
CCMA: Cognitive Control of Mobility in Aging
AD: Alzheimer’s Diseases
D-KEFS: Delis-Kaplan Executive Function System
IC: Independent Component
ERSPs: Event-related spectral perturbations
ANOVA: Analysis of Variance
SWV: Step width variability
SW: step width
FDR: False Discovery Rate
LSM: Left Sensorimotor
RSM: Right Sensorimotor
SD: Standard Deviation
BMI: Body Mass Index
LLFDI: Late Life Function and Disability Instrument
IADL: Independent Activities of Daily Living
LLFDI: Late-Life Function and Disability Instrument
RBANS: Repeatable Battery for the Assessment of Neuropsychological Status

## Acknowledgements

We like to acknowledge the mayor contributions by Dr. Joe Verghese and his team at the Department of Cognitive and Motor Aging. Regular weekly meetings with Dr. Verghese and frequent lab presentations were instrumental in shaping the ideas and hypothesis of this work. Finally, sincere thanks go to our participants for their willingness to volunteer for this research.

## Funding

The primary source of funding for this work was provided by a Mentored Research Scientist Development Award from the National Institute on Aging (K01AG049991). Participant recruitment was assisted by Dr. Jeannette Mahoney and supported by the National Institute on Aging (R01AG044007, R01AG036921, and K01AG049813) and by the Resnick Gerontology Center of the Albert Einstein College of Medicine. Participant evaluation and neuropsychological assessments were aided by the Human Clinical Phenotyping Core at Einstein, a facility of the Rose F. Kennedy Intellectual and Developmental Disabilities Research Center (RFK-IDDRC) which is funded by a center grant from the Eunice Kennedy Shriver National Institute of Child Health & Human Development (U54 HD090260, P50 HD105352-01).

## Competing interests

The authors report no competing interests.

## Notes

### Competing Interest Statement

The authors have declared no competing interest.

